# Hypoglycemia Aggravated Cognitive Degeneration by activating Endothelial ZBP1-mediated PANoptosis in Type 2 Diabetes

**DOI:** 10.64898/2026.03.17.712288

**Authors:** Wenping Luo, Qian Xiao, Na Li

## Abstract

Recurrent hypoglycemia increases cognitive impairment in diabetes mellitus patients. Following cerebral neuron injury, endothelial cells provide morphological, metabolic and immune support to damaged neurons. We investigated the inflammatory mechanism involved in hippocampal neuron degeneration. Behavioral experiments, including the open field test (OFT) and the Morris water maze test, were performed to measure cognitive changes. Using a vascular ring experiment, we evaluated vasodilation of the carotid artery. ZBP1 expression was knocked down after transfection with small interfering RNA in a brain endothelial cell line (bEnd3). In this study, PANoptosis, a recently defined form of programmed cell death (PCD), was found to be increased by hypoglycemia in the hippocampus of type 2 diabetic mice *in vivo* and by low glucose in bEnd3 cells *in vitro*. ZBP1 knockdown decreased PANoptosis induced by low-glucose stimulation in high-glucose-cultivated bEnd3 cells. RNA transcriptomics sequencing revealed that AGE-RAGE signaling significantly changed after ZBP1 was knocked down in bEnd3 cells. Corresponding biochemical data confirmed that ZBP1 knockdown regulated the advanced glycation end products (AGEs)-Receptor for Advanced Glycation End Products (RAGE) axis in bEnd3 cells. We present the first evidence that hypoglycemia impaired cognition in mice with type 2 diabetes by activating brain endothelial ZBP1-mediated PANoptosis via the AGE-RAGE axis.

**ARTICLE HIGHLIGHTS:**

- PANoptosis, a newly defined form of programmed cell death, is induced in the hippocampus after recurrent hypoglycemia in male db/db mice.
- ZBP1, a sensor of the PANoptosome, was activated in low glucose cultured brain endothelial cells.
- Hypoglycemia impairs vasodilation and cognitive function in type 2 diabetic mice.
- Our study indicates that inhibiting ZBP1-PANoptosis and the AGE-RAGE axis may be a potential approach to prevent hypoglycemia-induced cognitive degeneration in individuals with type 2 diabetes.

## 1. INTRODUCTION

The harmful effects of hypoglycemia on the course of type 1 [1] and type 2 diabetes [2] are multifaceted, especially the irreversible damage to the brain and cognitive function [3]. There are multiple causes of hypoglycemia, but the main causes include improper insulin injection [4] or glucose metabolism disorders after surgery [5]. Although mild or moderate hypoglycemia is not life-threatening, it can cause cognitive impairment if it occurs repeatedly [6]. The brain is extremely sensitive to fluctuations in blood glucose. Studies have indicated that recurrent hypoglycemia (RH) leads to cognitive impairment in diabetic patients [7] or diabetic mice [8]; however, the underlying mechanisms remain unclear.

Endothelial cells are the most important cells for maintaining the morphology and functions of blood vessels. Affected by blood glucose fluctuations [9], endothelial cells secrete various signaling factors to participate in the processes of nerve signal transmission, neuronal activity and other types of humoral regulation [10]. Activated c-Fos/focal adhesion preserved endothelial integrity and alleviated diabetic retinopathy [11]. c-Fos, a marker of neuronal activation, is involved in the processes of memory formation and learning and also regulates neural plasticity [12]. The hippocampus is a major organ involved in spatial learning and memory formation [13]. Won et al. (2012) reported that moderate hypoglycemia caused hippocampal dendrite damage, microglial activation, and cognitive impairment in rats with type 1 diabetes [14]. Most studies have focused on type 1 diabetes during episodes of hypoglycemia; however, type 2 diabetes accounts for a greater proportion of people with clinical diabetes. To provide more reliable evidence for better standardized blood glucose management and prevention of brain damage caused by hypoglycemia, it is necessary to understand the cognitive changes and related mechanisms in type 2 diabetes mellitus (T2DM) patients or mice during hypoglycemic episodes.

Inflammation is closely related to both energy metabolism in the brain and cognitive impairment [15]. The authors reported that hypoglycemia causes widespread inflammation throughout the body and may lead to serious adverse consequences such as cognitive impairment and dementia. The underlying molecular mechanism remains to be studied. PANoptosis is a new type of CD revealed in recent years and refers to a unique inflammatory cell death modality involving interactions among pyroptosis, apoptosis, and necroptosis. PANoptosis can be mediated by multifaceted PANoptosome complexes assembled by integrating components from other cell death modalities [16]. Z DNA binding protein (ZBP1) has been characterized as a critical innate immune sensor of not only viral RNA products but also endogenous nucleic acid ligands. ZBP1 sensing of the Z-RNA produced during influenza virus infection induces cell death in the form of pyroptosis, apoptosis, and necroptosis (PANoptosis) [17, 18]. The functions of these molecules in innate immunity and inflammatory cell death suggest new therapeutic targets for ZBP1-mediated diseases. A previous study revealed the progression of PANoptosis in diabetes mellitus and complications from diabetes [19], including cardiovascular diseases [20] and diabetic kidney injury [21]. However, whether PANoptosomes form in the context of hippocampal injury induced by hypoglycemia is unclear. In this study, the effects of hypoglycemia on mice with type 2 diabetes were assessed *in vivo* and *in vitro*, and we hope that these findings provide a promising landscape for studying diabetes treatment.

## 2. RESEARCH DESIGN AND METHODS

### 2.1 Animals

Male 16-week-old db/db (C57BL/KsJ) mice weighing 50–60 g was acquired from the Experimental Animal Center of Chongqing Medical University. They were stored in a 12 h light–dark cycle at 22°C–25°C with free access to food and water. The standard diet (SPF-F01-001, SpfBiotech, China) provided adequate nutrition with approximate compositions of ∼18-22% protein, ∼5-6% fat, and ∼4-5% fiber, with the remainder consisting of complex carbohydrates, vitamins, and minerals. Db/db mice were randomly assigned to the two groups (n = 6): the DM group and the DM recurrent hypoglycemia group (RH-DM). The RH-DM group indicates that hypoglycemia occurs in mice for 1 h per day and lasted for 5 days [22], which differentiated with the DM group due to its sharply fluctuated blood glucose. The Laboratory Animal Management Committee of Chongqing Medical University approved this study. The animal treatment method fully complied with the relevant guidelines of the Animal Protection Law of the People’s Republic of China and obtained approval from the Institutional Ethics Committee of Chongqing Medical University of the State Science and Technology Commission of the People’s Republic of China (Approval number: SC14,15-DHETK Chongqing 2007-0001).

### 2.2 Animal hypoglycemia models

To provoke a severe form of hypoglycemia, T2DM mice underwent subcutaneous morning injections of insulin (Wanbang, China) with 10.0 units/kg at 8:00–9:00 after overnight fasting[22]. The control group was administered the same volume of phosphate-buffered saline (PBS) instead of insulin to induce a transitory hypoglycemia state (hypoglycemia for 1 h/5 days). After insulin was injected, the blood samples were collected by a tail prick for blood glucose control every 30 min to guarantee the maintenance of glucose levels in severe hypoglycemia (<2.3mmol/L)[22]. Mice were injected with 50% glucose in PBS for hypoglycemia episode termination. None of them had seizures or coma during a hypoglycemia attack. The animals were fasted overnight (for about 12 hours) on the last day of experimental model before the induction of anesthesia or the collection of blood samples. All of them were sacrificed by sodium pentobarbital anesthesia (200 mg/kg)[23], and blood samples were collected by cutting the carotid artery. In addition,brain tissues were gathered for subsequent examinations.

### 2.3 Open field test

Before the start of the experiment, ensure that the laboratory box is clean and odor free, paying special attention to cleaning the stool and urine left by the animals in the previous experiment at the bottom of the laboratory box. The researchers should set appropriate parameters in the software, and record the serial number of animals, such as date and status information. Gently remove from its breeding of experimental animals, pay attention to make it back to the experimenter. Placed experimental animals quickly in the central area of the experiment box, and leave immediately. Open the animal behavior analysis software, record the activities of the animals in the casing, usually experiment time for 15 minutes. After the experiment, the experimental animals into the prepared other raise cage. Use of alcohol spraying apparatus of deodorant, dry with paper towel. The division of the area usually includes four corners, four sides and a central area. The software can automatically calculate the activity distance, stay time, entry times and average speed of experimental animals in these areas.

### 2.4 Morris water maze experiment

There are three phases in the experiment. Adaptation period: Let the animals swim freely in the pool and get familiar with the water maze environment. Learning stage: Put the animal into the water maze, let it find and remember the location of the platform, record the time (incubation period) and track of the animal to find the platform. Test phase: Remove the platform, let the animals swim in the pool for a certain period of time, and record the time and trajectory of the animals near the previous platform location.

### 2.5 Vascular ring experiment

Pre-cooling and oxygenation: The PSS solution is pre-cooled in a refrigerator at 4 ° C, then removed and oxygenated at a low flow rate with a mixture of 95% oxygen and 5% carbon dioxide for 20 minutes. Vascular ring preparation: The carotid arteries of the mice were removed immediately after execution and placed in an already cold and fully oxygenated PSS solution. The periaortic connective tissue was removed and about 2.5 mm carotid artery was intercepted, which was then placed in a cold, oxygenated PSS solution for temporary storage. Vascular balance: Add 5mL of pre-cooled PSS solution to each channel of the vascular tensiometer, and heat the remaining PSS in a 37℃ water bath. Then, the blood vessel segment prepared in step (2) is carefully placed on two steel needles, and the mixture is continued while the temperature is slowly raised to 37C. Every time thereafter, PSS or KPSS at 37 °C will be replaced. Adjust the vascular tension to about 4 mN, check every 5 minutes, if the tension drops back to 4 mN, and replace the PSS every 20 minutes. The vascular ring is in equilibrium in this state for 60 minutes. Until the vascular tone shows stability, at which point the system is zeroed out. Vascular activation: The PSS solution in the channel chamber was emptied, and the KPSS solution was replaced for stimulation for 15 minutes. The blood vessels contracted under the stimulation of KPSS, and the curve rose rapidly and gradually leveled off. After that, wash twice with PSS solution for 5 minutes each time. Add KPSS again for stimulation to ensure that the difference between the two contractions 15 minutes before and after is less than 10%. Otherwise, a third KPSS stimulus is performed until the requirement is met. After that, the PSS solution was used for three washes for 5, 10 and 15 minutes. Acetylcholine test: After the last wash, replace with a new 5mL PSS solution and perform systematic zeroing. Add 2μM of norepinephrine, and gradually add acetylcholine (concentration range from 10^-7^M to 10^-3^M) after the vascular tension is stabilized, observe and record the tension change. Sodium nitroprusside experiment: The blood vessels were washed five times with PSS solution for 5 minutes each time, after which 5 mL of new PSS solution was replaced and the system was zeroed again. 2μM of norepinephrine was added, and after the vascular tension reached a plateau, sodium nitroprusside was added successively (the concentration range was 10^-7^M to 10^-3^M), and the change of tension was recorded.

### 2.6 Immunofluorescent staining

Paraffin sections dewaxed to water. Antigen repair: Tissue sections were placed in a repair box filled with EDTA antigen repair buffer (PH9.0) in a microwave oven for antigen repair. Fire for 3min, ceasefire for 10min, turn to low fire for 3min, this process should prevent excessive evaporation of buffer, do not dry. After natural cooling, the slide was placed in PBS (PH7.4) and washed by shaking on the decolorizing shaker for 3 times, 5min each time. Blocking: Paraffin sections were inserted into PBS for 3min*3 times, and each section was sealed with 200ul 5% Bovine Serum Albumin (BSA) for 30min. Incubation of primary antibody: Dilute primary antibody (c-FOS, ABclonal, A24620) with PBS to an appropriate concentration, add 50ul of primary antibody to each tablet, and incubate overnight at 4℃. Incubation of secondary antibodies: the slices were taken out of the refrigerator and rewarmed for 20-30min, washed with PBST for 3 times, diluted the FITC-labeled secondary antibodies with 1% BSA, added 50ul of secondary antibodies to each slice, and incubated in the oven at 37℃ for half an hour. Redyeing DAPI: PBST was washed for 3min*3 times, diluted DAPI was added and incubated for 10min at room temperature. Microscopy: Each section drops an appropriate amount of anti-fluorescence quench agent to cover the cover glass, fluorescence microscope observation.

### 2.7 Hematoxylin and eosin (H&E) staining

Paraffin sections dewaxed to water. Hematoxylin, 3 min, then wash with distilled water to remove floating color. 1% hydrochloric acid alcohol differentiation solution. Middle differentiation 30s→ tap water I, 3min → tap water II, 3min. Eosin dye solution, 2 min, then wash with distilled water to remove floating color. Dehydration: 75% ethanol, 85% ethanol, 95% ethanol, 100% ethanol for 1 min each. Transparent: Xylene I, 1 min→ xylene II, 1 min. Sealing and scanning.

### 2.8 Cell culture and treatment

Brain endothelial cells (bEnd3, TCM-C715, HyCyte, China) was cultivated at 37 °C with 5% CO_2_ in the bEnd.3 cell-specific culture medium (TCM-G715, Hycyte, China), which was completed medium that supplemented with 10% (v/v) FBS and 1% (v/v) penicillin/streptomycin (PS). High-glucose treatment (30 mmol/L) was lasted for 72 hours. Low-glucose medium was composed of Dulbecco ’ s Modi ed Eagle Medium (DMEM) (Invitrogen, Carlsbad, CA, USA) contain 1 mM glucose supplemented with 10% (v/v) FBS (Invitrogen) and 1% (v/v) penicillin/streptomycin (PS) (Beyotime Biotechnology, Jiangsu, China). The components of the basic culture medium are the same, but the difference lies in the concentration of glucose. Based on the published study methods, cells were treated with high-glucose cell culture medium, which contains 30 mM glucose for 3 days or low-glucose cell culture medium, which contains 1 mM glucose for 1 h, and then replaced with high-glucose medium for another 1 h. This step was repeated three times to modulate the process of recurrent hypoglycemia[24].

### 2.9 Small interfering RNA transfection

Bend.3 cells were incubated in the six-well plate 1 day before transfection. The second day, cells were transfected with 50nM mouse siRNA or negative control according to the manufacturer’s guidelines (RM09014P, ABclonal, China). Cells were added to each well with 8 μL transfection reagent. After 6 hours, cultured medium was half-replaced with fresh completed medium. After 48 hours, total protein was extracted using Western blot analysis. The siRNA sequences for AIM2 were 5′-GCAGUGACAAUGACUUUAATT-3′and 3′ -UUAAAGUCAUUGUCACUGCTT-5′. The siRNA sequences for RIPK1 were 5′-GGCAGAAUGAGGCUUACAATT-3′ and 3′-UUGUAAGCCUCAUUCUGCCTT-5′. The siRNA sequences for ZBP1 were 5′-GAGACAAUCUGGAGCAAAATT3′and 3′-UUUUGCUCCAGAUUGUCUCTT-5′, respectively. The siRNA sequences for RAGE were 5′-ACCGGGGGCATTCAGCT-3′ and 3′-TCTAAAAAAGGGCATTCAGCTGT-5′. The negative control sequences for those small interfering RNA were sense strand: 5’-UUC UCC GAA CGU GUC ACG UTT-3’ and antisense strand: 5’-ACG UGA CAC GUU CGG AGA ATT-3’.

### 2.10 RNA sequencing

Total RNA was extracted from bEnd.3 transfected with siNC or siZBP1 after HG+LG treatment using the TRIzol reagent (Invitrogen) according to the manufacturer’s instructions. A total amount of 1 μg RNA was used to prepare the sequencing libraries using an NEBNext Ultra RNA Library Prep Kit for Illumina (NEB, Ipswich, MA) following the manufacturer’s recommendations. The qualified libraries were sequenced on an Illumina NovaSeq 6000 platform to generate 150 bp paired-end reads by a commercial service provider, like Novogene/Gene Denovo. Differential gene expression analysis between siNC and siZBP1 groups was performed using the DESeq2 or edgeR package in R, with genes showing a |log2(Fold Change)| > X and an adjusted p-value (FDR) < 0.05 considered statistically significant. Gene Ontology (GO) enrichment and Kyoto Encyclopedia of Genes and Genomes (KEGG) pathway analysis of the differentially expressed genes (DEGs) were conducted using the clusterProfiler package in R, with an adjusted p-value < 0.05 considered significant.

### 2.11 Flow cytometry

The gathered cells were rinsed with PBS and 2% fetal calf serum. Each examination was performed in non-dyed, annexin V, PI, and PI and annexin V double-dyed groups. The treatment group underwent double dyeing from low to high. The 4× binding buffer was diluted to 1× buffer with PBS, and after the residual PBS in the centrifuge tube was absorbed, 100 μL of 1× binding buffer was added to each tube, and the cells were blown with a pipette to fully resuspend the cells, whereas the dyes were added under the condition of dark light. Annexin V or PI 5 μL was added to the single-dyed and double-dyed groups and gently mixed with a pipette. After the incubation at room temperature and darkness for 15 min and mixing with 1×binding buffer 300 μL, the cell suspension was removed to a 5 mL flow tube under dark light and was analyzed by flow cytometry within 1 h. Annexin V-FITC/PI apoptosis detection kit was used as a probe with annexin V annexin V-labeled FITC. The maximum excitation wavelength of FITC was 488 nm, the maximum emission wavelength was 525 nm, and the green fluorescence of FITC was found in the FL1 channel. The maximum excitation wavelength of the PI–DNA complex comprised 535 nm, the maximum emission wavelength was 615 nm, and the red fluorescence of PI was outlined in the FL2 or FL3 channel. A double dispersion plot was constructed through software analysis, with FITC as the horizontal coordinate and PI as the vertical.

### 2.12 Western blotting

Hippocampus tissues were placed in liquid nitrogen, turned into powder, and then removed to a 1.5 mL centrifuge tube. Subsequently, four volumes (μL/mg) of lysis buffer (1% Triton X-100, 1% protease inhibitor, and 1% phosphatase inhibitor) were complemented by powder and incubated for 1 h on Rotater at 4°C. The cultivated cells were immersed in lysis buffer for 30 min. Supernatants were obtained by centrifuge (12,000 ×g × 15 min at 4°C). Subsequently, protein concentration was calculated with a BCA protein assay kit based on the manufacturer’s instructions (23225, Thermo Fisher Scientific). Protein samples (20 µg/lane) underwent electrophoresis in a 4%–20% SDS polyacrylamide gel (P0468S and P0469S, Beyotime) and mounted onto polyvinylidene fluoride membranes (FFP39, Beyotime) fixed with 5% milk in TBST for 1.5 h and infested with primary antibodies (NLRP3, Proteintech, 68102-1-Ig; PYRIN, Abcam, ab195975; RIPK1, Proteintech, 29932-1-AP; RIPK3, Proteintech, 17563-1-AP; Casepase-8, Proteintech, 13423-1-AP; Caspase-1, Proteintech, 22915-1-AP; ZBP1, Proteintech, 13285-1-AP;AIM2, Proteintech, 20590-1-AP; N-GSDMD, Proteintech, ab215203; GSDMD, Proteintech, 20770-1-AP; MLKL, Proteintech, 66675-1-Ig; Caspase-3, Proteintech, 66470-2-Ig; FADD, Proteintech, 14906-1-AP;ASC, Proteintech, 10500-1-AP; IL-18, Proteintech, 10663-1-AP; IL-1β, Proteintech, ab283818; Casepse-9, Proteintech, 66169-1-Ig; Bcl2, Proteintech, 12789-1-AP; BAX, Proteintech, 50599-2-Ig; β-actin, Servicebio, GB15001-100) overnight at 4°C. The membranes were rinsed thrice with TBST and then with HRP-labeled species-specific secondary antibodies (HRP conjugated Goat Anti-Rabbit IgG (H+L), Servicebio, GB23303; HRP conjugated Goat Anti-Mouse IgG (H+L), Servicebio, GB23301) for 1 h. Subsequently, the membranes were washed thrice with TBST. Blotting with BeyoECL HRP substrate (P0018AM, Beyotime) was performed in a ChemiDoc XRS (Bio-Rad) image acquisition system. Quantitative densitometry evaluation of all bands was carried out with Image J processing software.

### 2.13 Cell viability assay

The cell viability was detected by Cell Counting Kit-8 (CCK8, HY-K0301, Med Chem Express, USA) assay. Brain endothelial cells (bEnd3, TCM-C715, HyCyte, China) was seeded in 96-well growth medium plate overnight at 1*10^4^ cells/well. After 24 h, the medium was replaced, and cells were stimulated with low-glucose medium. Next, cells were maintained at 37℃ and 5% CO_2_ in a humidified incubator for different hours, then these cells were cultured with CCK-8 for 30 min.

### 2.14 Statistical analysis

All the data were statistically analyzed using SPSS (version 22.0; IBM, Armonk, NY, USA). Statistical analyses were performed using GraphPad Prism version 8.0.2 (GraphPad Software, Inc.). Independent samples t tests were used for comparisons between two groups, and one-way ANOVA with Tukey’s multiple comparison test for multiple group comparisons. All data were shown as mean ± SEM. A *P* < 0.05 was regarded as statistically significant. Data from at least three independent experiments were evaluated. [25].

## 3. RESULTS

### 3.1 Recurrent hypoglycemia impaired the cognition and vasodilation of mice with type 2 diabetes

We established a reliable model for recurrent hypoglycemia in db/db mice, a model for type 2 diabetes. The blood glucose (BG) levels of the type 2 diabetes mellitus (DM) group and the recurrent hypoglycemia-diabetes mellitus (RH-DM) group are shown in Figure 1a and Figure 1b, respectively. The blood glucose concentration of the DM group was consistently higher than 20 mmol/L, much higher than the criteria for a diabetes diagnosis (random BG concentration ≥11.1 mmol/L). However, after subcutaneous injection of insulin, the blood glucose concentration of the mice in the RH-DM group was reduced to less than 3 mmol/L and maintained at that level for 1 hour. The process was repeated every day for five days.

**Figure 1.**
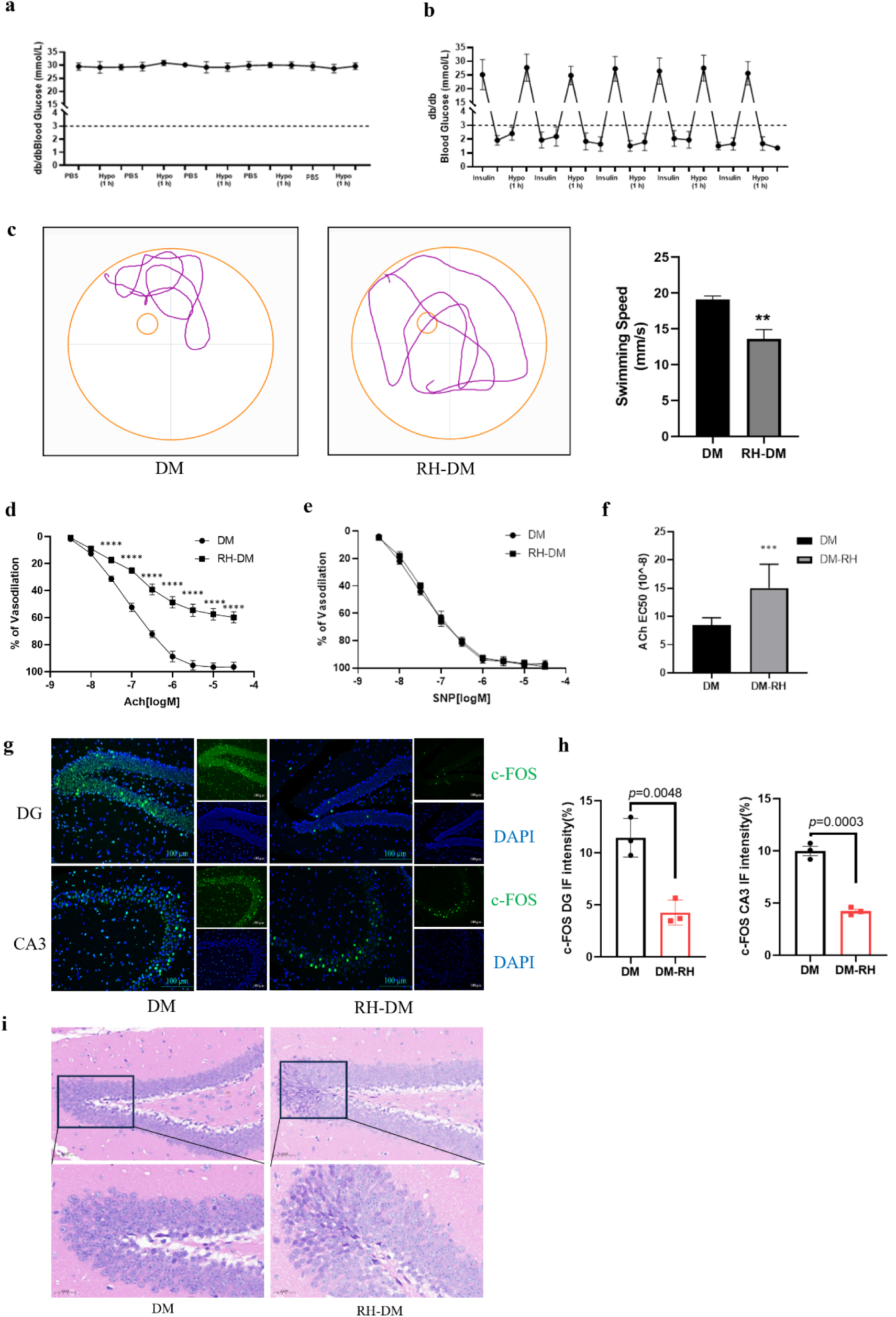
Recurrent hypoglycemia damaged cognition and vasodilation of type2 diabetic mice. **a**, Blood glucose fluctuation curve of type 2 diabetic db/db mice (DM group, n=6). **b**, Blood glucose fluctuation curve of recurrent hypoglycemia-type 2 diabetic db/db mice (RH-DM group, n=6). **c**, Representative images of Morris water maze traces and the quantitative swimming speed of mouse in the DM and the RH-DM group. **d**, Carotid endothelium dependent vasodilation was tested by the acetylcholine test (n=6). **e**, Carotid endothelium dependent vasodilation was tested by the Sodium nitroprusside experiment (n=6). **f**, AchE50 statistical analysis of DM and RH-DM group mice carotid artery (n=6). *** *p*<0.001 vs DM group; **** *p*<0.0001 vs DM group. **g-h**, Immunofluorescent staining and statistical analysis of c-FOS (a marker of neuronal activity in neuroscience) in hippocampus of the DM group and the RH-DM group. **i**, H&E staining of mouse brain in the DM and the RH-DM group. AchE50 means the concentration of the compound required to reduce acetylcholinesterase activity by 50%. Higher AchE50 value indicates lower inhibitory effects and lower acetylcholinesterase activity.DM: diabetes mellitus; RH-DM: Recurrent hypoglycemia -diabetes mellitus.

The effects of hypoglycemia on spatial learning and cognitive function were evaluated by open field tests (OFTs) and Morris water maze experiments. Table 1 shows the OFT results, in which spontaneous activity, exploratory behavior, anxiety and depression were quantitatively evaluated. The results indicated that the central lattice times of the DM and RH-DM groups were 33.48±16.61 s and 19.25±11.95 s, respectively (*p*<0.01). The horizontal scores of the two groups were 6.8±2.5 and 3.3±1.7 (*p*<0.01), respectively, and the vertical scores were 0.5±0.3 and 0.2±0.1 (*p*<0.01), respectively. The grooming and urinating times in the RH-DM group were 1.2±0.5 and 0.3±0.2, respectively, which were lower than those in the DM group.

**Table 1.** The effects of hypoglycemia on spatial learning and memory function in diabetic mice were evaluated by open field experiment.

| Index | DM ( n=6 ) | RH-DM ( n=6 ) | T value | P value |
| --- | --- | --- | --- | --- |
| Central lattice time(s) | 33.48±16.61 | 19.25±11.95 | 2.979 | **0.0074** |
| Horizontal score | 6.8±2.5 | 3.3±1.7 | 3.896 | **0.0009** |
| Vertical score | 0.5±0.3 | 0.2±0.1 | 3.267 | **0.0039** |
| Grooming(time) | 2.7±1.3 | 1.2±0.5 | 3.697 | **0.0014** |
| Urinate(time) | 0.6±0.4 | 0.3±0.2 | 2.285 | **0.0333** |

The maze method, especially the water maze method (Table 2), uses an animal’s instinctive response to escape from drowning, and the behavioral model is a model of spatial memory related to the hippocampus [26]. In this study, mice were trained in a water maze to learn to distinguish spatial orientation to provide a quantitative measure of behavioral phenotypes for each group. Compared with the DM group, the platform crossing times was 4.4±0.84, and the platform quadrant residence time was 10.54±1.71 (s); these indices were reduced to 1.23±1.03 and 6.22±1.09 (s), respectively for the RH-DM group (n=6; *p*<0.0001). The swimming distance traveled in the platform quadrant in the RH-DM group was reduced compared to the DM group (n=6; *p*<0.0001). A reduction in the intensity and frequency of activity revealed that hypoglycemia suppressed the excitability of diabetic mice; in other words, it increased their depression. Representative images of the water maze experiment are shown in Figure 1c, and the swimming trajectory of the RH-DM group was more complicated than that of the DM group. Compared with those in the DM group, the crossing times and residence times in the other three quadrants in the RH-DM group clearly increased. Moreover, the reduced frequency of crossing the original platform quadrant in the RH-DM group indicated that hypoglycemia impaired the spatial learning and memory ability of the mice.

**Table 2.**
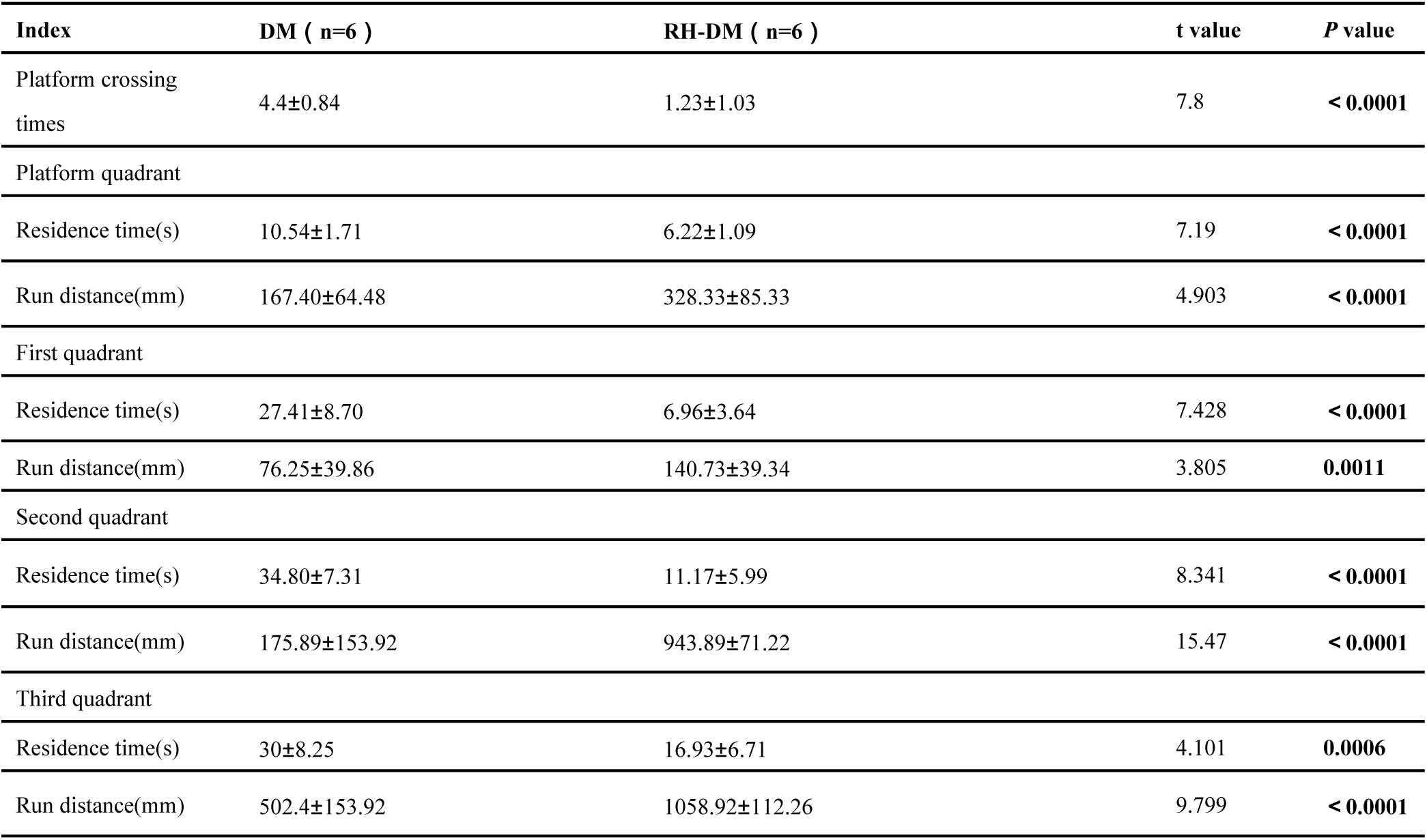
The effects of hypoglycemia on spatial learning and memory function in diabetic mice were evaluated by Morris water maze experiment.

Changes in vasodilation function are closely related to brain cognitive function [27]. In this study, a vascular tension test was carried out to evaluate the vasodilative function of the carotid artery. The percentage changes in vascular tension after the gradual addition of acetylcholine (the concentration ranged from 10^-7^ M to 10^-3^ M) are shown in Figure 1d. In the DM group, the percentage of vasodilatory tension decreased more sharply than that in the RH-DM group did (n=6; *p*<0.0001). However, the changes in the tension percentage after the addition of sodium nitroprusside (at concentrations ranging from 10^-7^ M to 10^-3^ M) did not significantly differ between the two groups ( Figure 1e). The mean concentrations for 50% of the maximal effect of Ach (Ach EC50) in the DM and RH-DM groups were 8.446×10^-8^ M and 15.03×10^-8^ M (Figure 1f, *p*<0.001), respectively.

The hippocampus plays a critical role in cognition, spatial learning and memory [28]. We conducted immunofluorescence staining of c-FOS (Figure 1g), which is a molecular marker of neuronal activity and memory formation [12]. Compared with those in the DM group, the density of c-FOS immunofluorescence in the DG area and CA3 area in the hippocampus in the RH-DM group sharply decreased (Figure 1h). Additionally, H&E staining was conducted to assess inflammatory infiltration in the hippocampus. As shown in Figure 1i, acute inflammation was increased in the RH-DM group compared with the DM group.

### 3.2 PANoptosis was significantly induced by recurrent hypoglycemia in the diabetic mouse hippocampus *in vivo* and in endothelial cells *in vitro*

The mechanism of hippocampal cognitive dysfunction induced by hypoglycemia in mice with diabetes is still unclear. However, inflammation was found to occur widely [29]; thus, we tested whether different types of PCD were induced in the hippocampi of DM and RH-DM mice (Figure 2a–d). Receptors of PANoptosomes, including Receptor-interacting protein kinase 1 (RIPK1), Z-DNA binding protein 1 (ZBP1) and Absent In Melanoma 2 (AMI2), were activated in RH-DM mouse hippocampus compared with that in the DM group, respectively (Figure 2a, n=6; p<0.0001). The expression of the pyroptosis markers Nucleotide-binding oligomerization domain, leucine-rich repeat and pyrin domain-containing 3 (NLRP3), N-terminal of Gasdermin D (GSDMD), and Fas-associating via death domain (FADD) was increased in the hippocampus of the RH-DM group compared with that in the DM group (Figure2b, *n*=6; *p*<0.0001). The expression of apoptosis markers, including BAX, cleaved-caspase3, caspase-3 and caspase-8, was elevated in the RH-DM mouse hippocampus by approximately 1.8-fold compared with that in the DM group (Figure 2c, *n*=6; *p*<0.0001). Moreover, the expression of necroptosis markers, including phosphorylated and total Receptor-interacting protein kinase 3 (RIPK3) and Mixed lineage kinase domain-like protein (MLKL), obviously increased in the RH-DM group compared with the DM group (Figure 2d, *n*=6; *p*<0.0001). Representative immunofluorescence images of Caspase1, Caspase3 and RIPK3 expression in the mouse brain are shown in Figure 2c. The immunofluorescence intensity of these molecules increased in the RH-DM mouse hippocampus compared with that in the DM group (Figure 2e). Briefly, PANoptosis, including pyroptosis, apoptosis and necroptosis, was strongly increased in the hippocampus of hypoglycemia-stimulated diabetic mice.

**Figure 2.**
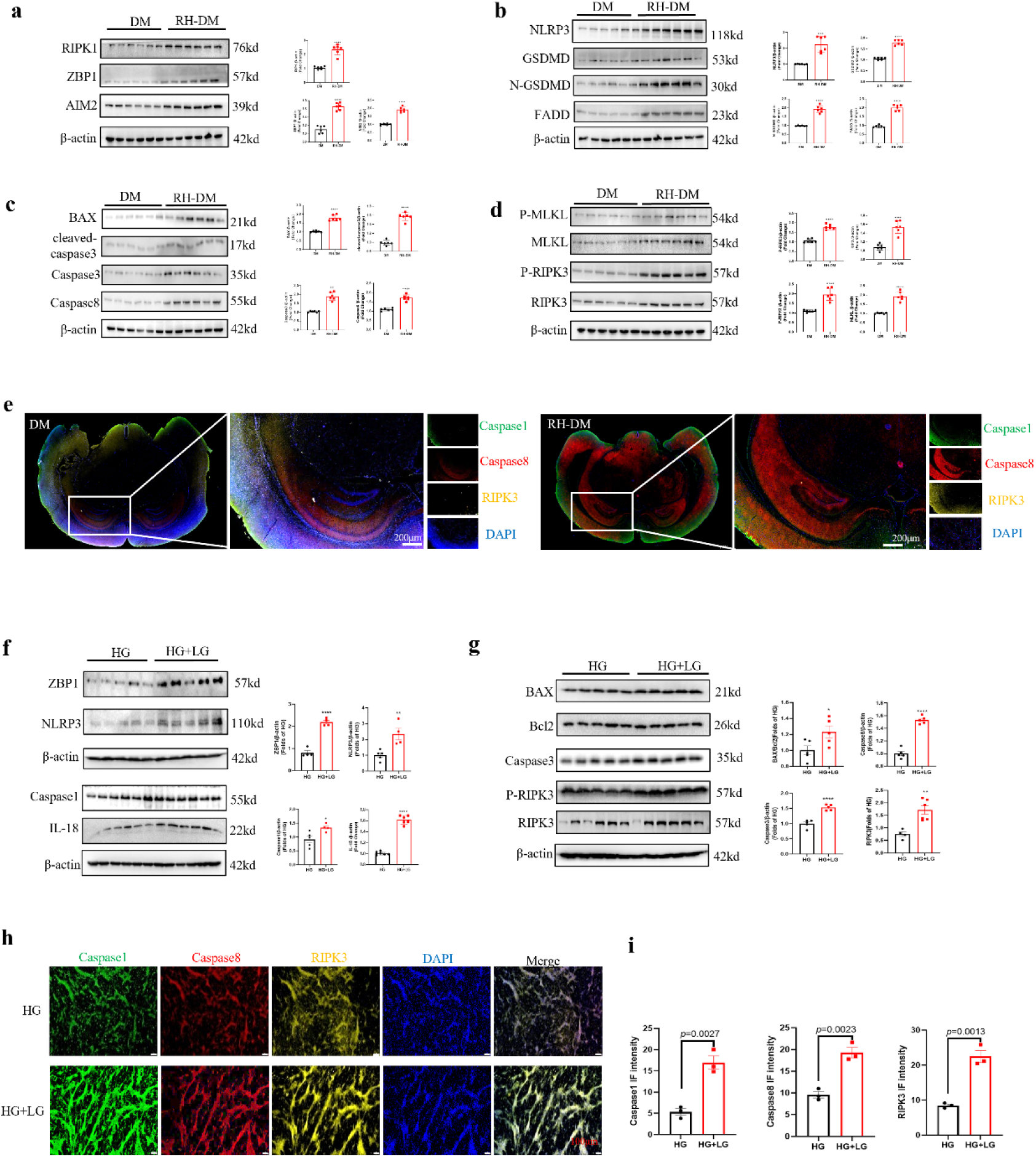
Hypoglycemia exacerbated PANoptosis in type 2 diabetic hippocampus *in vivo* and in bEnd.3 *in vitro*. **a**, The western blotting and statistical analysis of receptors of PANoptosme in hippocampus of type 2 diabetic mice (*n*=6). **b**, The western blotting and statistical of pyroptosis-associated markers in hippocampus of type 2 diabetic mice (*n*=6). **c**, The western blotting and statistical of apoptosis-associated markers in hippocampus of type 2 diabetic mice (*n*=6). **d**, The western blotting and statistical of necroptosis-associated markers in hippocampus of type 2 diabetic mice (*n*=6). \*\**p*<0.01 vs DM group; *** *p*<0.001 vs DM group; **** *p*<0.0001 vs DM group. **e**, Immunofluorescence staining of Caspase1, Caspase8 and RIPK3 in the hippocampus of the DM and RH-DM mouse. **f-g**, Western blotting of pyroptosis, apoptosis and necroptosis markers in bEnd.3 treated by HG or HG+LG. **e**, Statistical analysis of images in Figure 2c (*n*=5-6). **h-i**, Immunofluorescence staining and statistical analysis of Caspase1, Caspase8 and RIPK3 in the bEnd.3 after HG or HG+LG treatment. \**p*<0.05 vs HG group; ** *p*<0.01 vs HG group, **** *p*<0.0001 vs HG group. HG: high glucose; LG: low glucose; RIPK3: receptor-interacting serine-threonine kinase 3.

Brain endothelial cells are sensitive to blood glucose fluctuations [30]. We detected the effect of low glucose in the culture medium on brain-derived endothelial cells.3 (bEnd.3). Similarly, the *in vitro* expression of the PANoptosis marker in bEnd.3 cells was detected by western blotting, as shown in Figure 2f–g. The expression of ZBP1 in the HG+LG group was more than 2-fold greater than that in the HG group (Figure 2f, *n*=5; *p*<0.0001). The expression of the pyroptosis-associated markers NLRP3, caspase1 and IL-18 significantly increased in the HG+LG group (Figure 2f, *n*=5; *p*<0.05). Markers of apoptosis and necroptosis, including BAX/Bcl2, caspase3 and P-RIPK3/RIPK3 were elevated in the HG+LG group compared with those in the HG group (Figure 2g, *n*=5; *p*<0.05). Compared with those in the HG group, the immunofluorescence intensity of Caspase1, Caspase8 and RIPK3 in the HG+LG group was increased (Figure 2f–g). In summary, PANoptosis was activated by low glucose in high glucose-cultured bEnd.3 cells.

### 3.3 ZBP1 was activated in PANoptosis induced by low glucose in bEnd.3 cells *in vitro*

To determine which sensor of the PANoptosome was activated in endothelial PANoptosis, we transfected small interfering RNA of three different sensors (ZBP1, AIM2 and RIPK1) into the bEnd.3 cell line.

As shown in Figure 3a, compared with those in the HG group, the protein levels of necroptosis markers (p-RIPK1, RIPK1, p-MLKL and MLKL) clearly increased in the HG+LG group. ZBP1 was successfully knockdown by transfected with siRNA in bEnd.3 (Supplementary Figure1a). However, ZBP1 knockdown decreased the expression of necroptosis-associated markers, including phosphorylated RIPK1, RIPK1, phosphorylated MLKL and MLKL (*n*=3; *p*= 0.0399). The levels of apoptosis-associated markers, including caspase8, caspase9 and caspase3, were greater in the HG+LG group than in the HG group (*n*=3; *p*=0.0072). In contrast, blocking ZBP1 sharply decreased the apoptotic index (*n*=3; *p*=0.0019). Similarly, after ZBP1 expression was knocked down, we detected pyroptosis-associated markers in the HG+LG group (*n*=3; *p*=0.0130). The expression of ASC, GSDMD and IL-18 was elevated by low glucose stimulation compared with that in the HG group (*n*=3; *p*=0.0075). However, the expression of these PANoptosis-associated markers decreased in response to ZBP1 knockdown (Figure 3b) (*n*=3; *p*=0.0040). Apoptosis was detected by Hoechst/PI staining (Figure 3c). Statistical analysis revealed that compared with HG, ZBP1 knockdown decreased the apoptosis induced by low glucose in bEnd.3 cells (Figure 3d). Moreover, compared with that in the HG group, ROS production in bEnd.3 cells was increased in the HG+LG group but was reversed after ZBP1 expression was knocked down (Figure 3e–f). The rate of apoptosis was determined by flow cytometry, as shown in Figure 3g–h. Apoptosis stimulated by low glucose was attenuated by silencing ZBP1.

**Figure 3.**
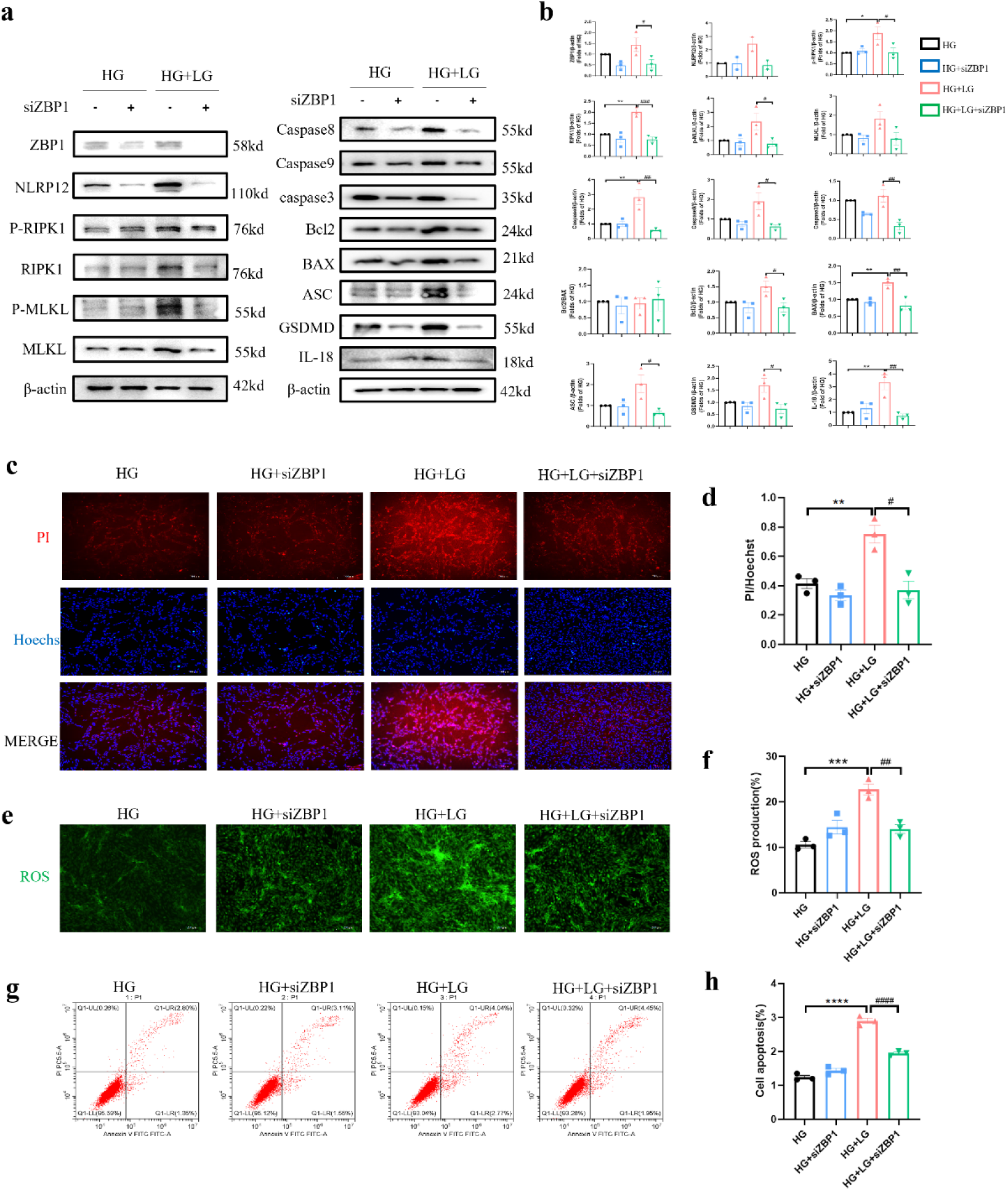
ZBP1 knockdown alleviated low glucose-induced PANoptosis and cell injury in bEnd.3. **a**, Western blotting of pyroptosis, apoptosis and necroptosis markers after knockdown ZBP1 expression in HG or HG+LG treated bEnd.3 *in vitro*. **b**, Statistical analysis of western blotting in Figure 3a. **c,** Hoechst/PI staining of bEnd.3 after ZBP1 knockdown treated with HG or HG+LG. **d,** Statistical analysis of PI/Hoechst ratio. **e-f,** Representative images and statistical analysis of ROS production in bEnd.3 after ZBP1 knockdown treated HG or HG+LG. **g-h**, Cell apoptosis tested by flow cytometry in bEnd.3 after knockdown ZBP1 treated with HG or HG+LG. \**p*<0.05 vs HG group, ** *p*<0.01 vs HG group, **** *p*<0.0001 vs HG group; ^#^*p*<0.05 vs HG+LG group, ^##^*p*<0.001 vs HG+LG group, ^###^*p*<0.001 vs HG+LG group. ZBP1: Z-DNA binding protein 1.

Interestingly, after silencing the expression of AIM2 (Supplementary Figure 1b; *n*=3; *p*=0.006) or RIPK1(Supplementary Figure1c; *n*=3; *p*=0.0008), there was no significant change in the expression of the PANoptosis-associated markers mentioned above (Figure 4a-d). For instance, the expression of ASC (*n*=3; *p*=0.6046), GSDMD-N/GSDMD (*n*=3; *p*=0.8705), BAX/Bcl2 (*p*=0.5964) and p-RIK3/RIPK3 (*p*=0.1771) changed little after knocking down AIM2 (Figure 4a-b). In addition, ASC (*n*=3; *p*=0.3603), GSDMD-N/GSDMD (*n*=3; *p*=0.2439), P-MLKL/MLKL (*n*=3; *p*=0.3797) and P-RIPK3/RIPK3 (*n*=3; *p*=0.7667) hardly changed after knocking down the expression of RIPK1, while the expression of cleaved-caspase8/caspase8 (*n*=3; *p*=0.007) and BAX/Bcl2 (*n*=3; *p*=0.0364) reduced obviously. In summary, ZBP1 was the sensor that mediated low glucose-induced PANoptosis in bEnd.3 cells. However, the mechanism by which ZBP1 mediates PANoptosis in the brain endothelium and hippocampus in the context of hypoglycemia needs to be clarified.

**Figure 4.**
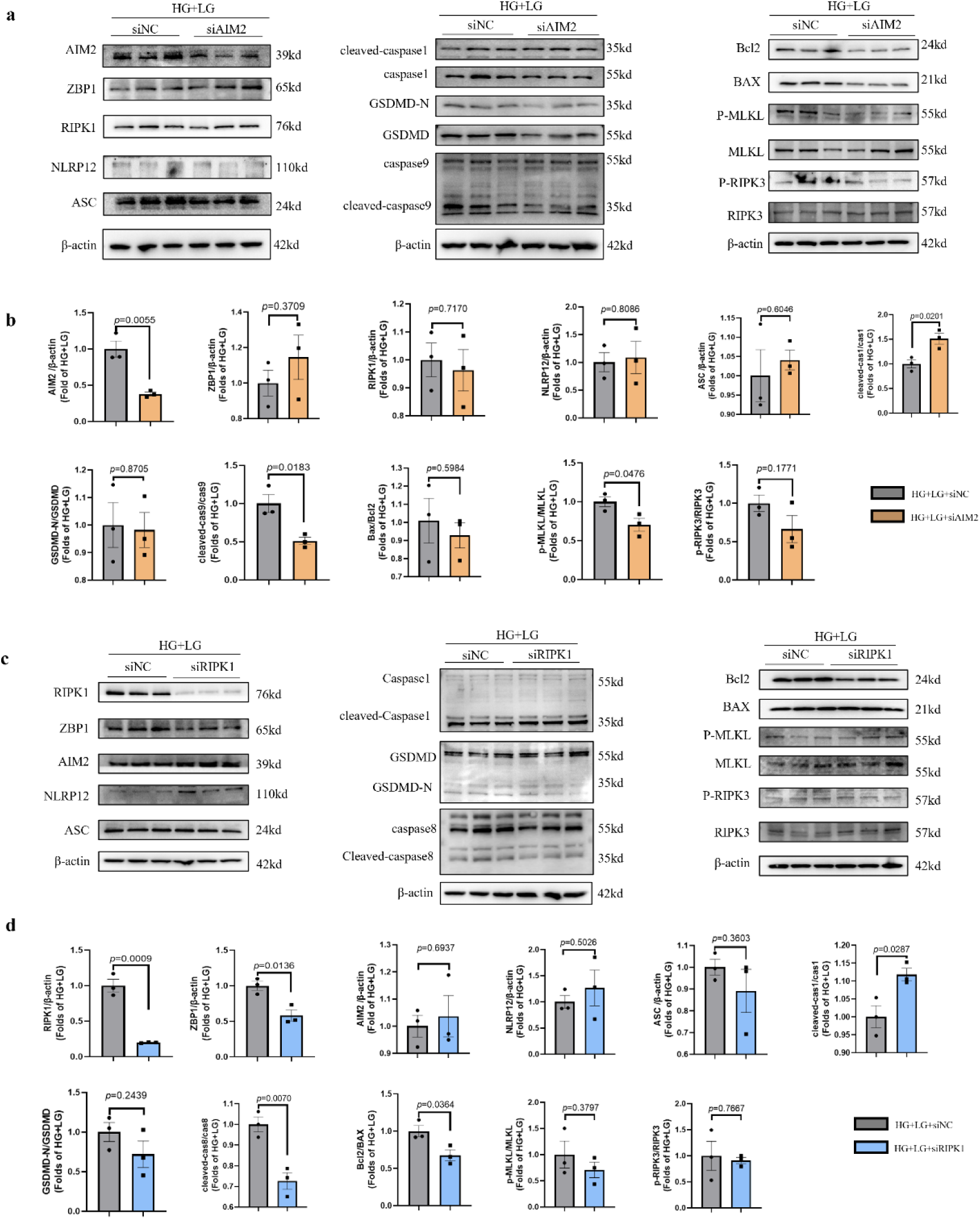
PANoptosis-markers has no significant changes after knocking down AIM2 or RIPK1 in HG or HG+LG treated bEnd.3 cell line. **a**, Western blotting of pyroptosis, apoptosis and necroptosis molecules in HG+LG treated bEnd.3 after knocking down the expression of AIM2. **b**, Statistical analysis of western blotting in Figure 4a. (*n*=3). **c**, Western blotting of pyroptosis, apoptosis and necroptosis molecules in HG+LG treated bEnd.3 after knocking down the expression of RIPK1. **d**, Statistical analysis of western blotting in Figure 4c. (*n*=3). AIM2: absent in melanoma 2; RIPK1: receptor interacting serine/threonine kinase 1.

### 3.4 Low glucose activated ZBP1-mediated PANoptosis via the AGEs/RAGE axis in vitro

We conducted RNA transcriptome sequencing in bEnd.3 cells after ZBP1 expression was knocked down. The volcano plot and heatmap of differentially expressed genes in bEnd.3 cells after ZBP1 expression was knocked down are shown in Figure 5a and Figure 5b, respectively. After ZBP1 was knocked down, 28937 genes were upregulated, and 89 genes were downregulated in bEnd.3 cells. KEGG enrichment analysis of DEGs in bEnd.3 cells revealed that the activity of the AGE-RAGE signaling pathway, which is closely related to diabetic complications, significantly changed (Figure 5c). Immunofluorescence staining revealed that hypoglycemia increased the intensity of RAGE in the hippocampal DG area in the RH-DM group compared with that in the DM group (Figure 5d). In addition, we detected the expression of RAGE after low-glucose treatment and si-ZBP1 transfection in bEnd.3 cells. Western blotting revealed that ZBP1 knockdown attenuated the expression of RAGE (Figure 5e–f) (n=3; *p*=0.0359). Moreover, immunofluorescence staining indicated that low glucose concentrations increased the intensity of RAGE expression on the cell membrane of bEnd.3 cells (Figure 5g), and ZBP1 knockdown reduced the immunofluorescence intensity of RAGE in bEnd.3 cells (Figure 5h) (n=3; *p*=0.0003). To determine the role of RAGE in ZBP1-mediated PANoptosis induced by HG+LG in bEnd.3 cells, we tested the expression of PANoptosomes after silencing RAGE in bEnd.3 cells (Supplementary Figure1d; Figure 5i–j). Statistical analysis revealed that RAGE knockdown reversed ZBP1-mediated PANoptosis induced by HG+LG in bEnd.3 cells. The results above suggested that ZBP1 regulated the AGE-RAGE axis, which could be the mechanism of low glucose-induced endothelial PANoptosis in the mouse hippocampus.

**Figure 5.**
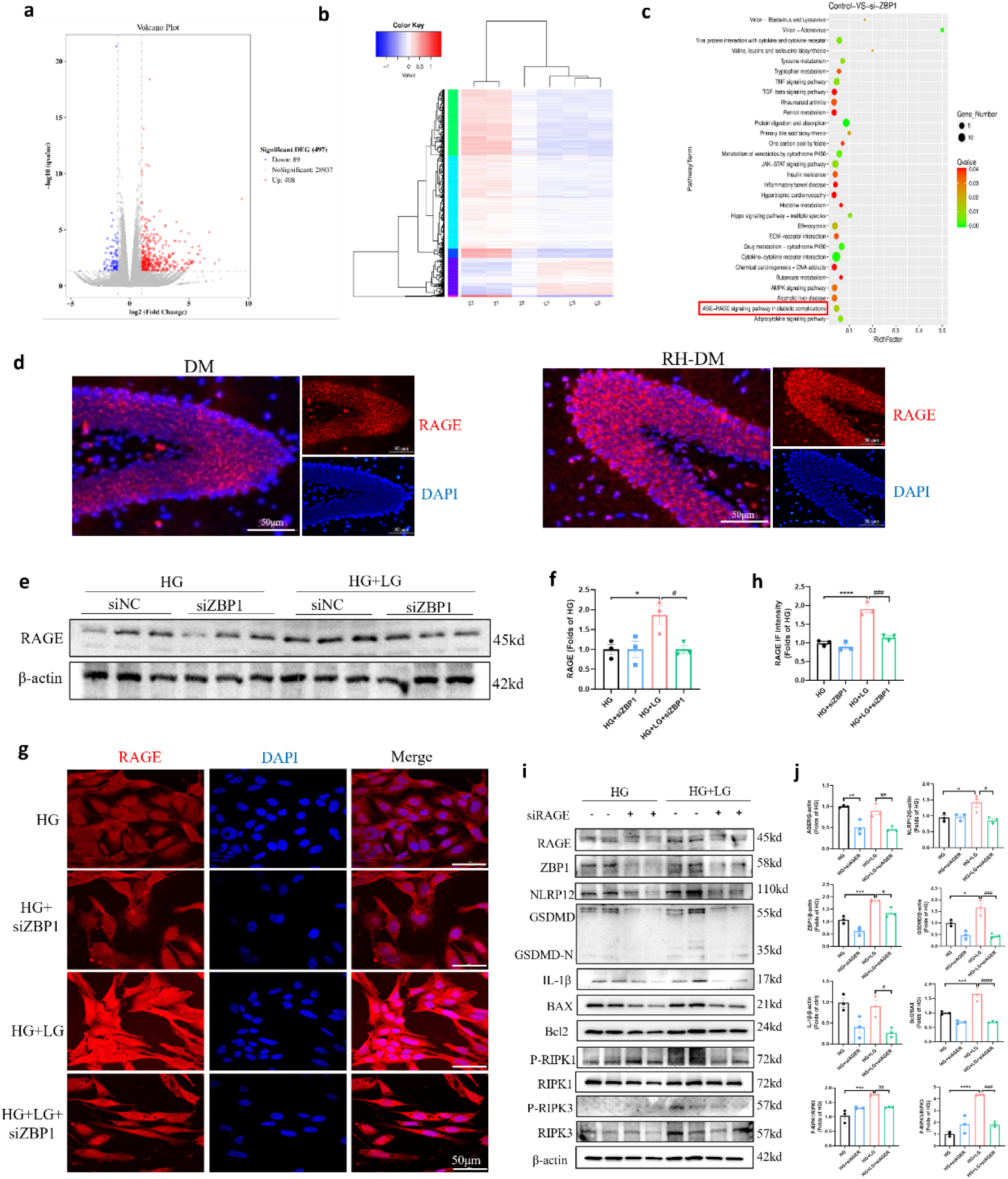
ZBP1-mediated PANoptosis was activated by hypoglycemia via AGEs-RAGE stress in bEnd.3. **a**, Volcano plot of differential expression genes in bEnd.3 after knocking down the expression of ZBP1. **b**, Heat map of differential expression genes in bEnd.3 after knocking down the expression of ZBP1. **c**, KEGG enrichment analysis bubble chart of differential genes in bEnd.3 after knocking down the expression of ZBP1. **d**, Immunofluorescent staining of RAGE in mouse hippocampus DG area of the DM and RH-DM group. **e**, Western blotting of RAGE after knocking down ZBP1 in HG or HG+LG treated bEnd.3. **f**, Statistical analysis of western blotting in figure 5e (*n*=3). **g**, Representative immunofluorescent staining images of RAGE in bEnd.3 after silencing ZBP1 expression in HG or HG+LG group. **h**, Statistical analysis of immunofluorescent density in figure 5g (*n*=3). **i,** Western blotting of RAGE and ZBP1-associated PANoptosis molecules after knocking down RAGE in HG or HG+LG treated bEnd.3. **j,** Statistical analysis of western blotting in figure 5i (*n*=3). \**p*<0.05 vs HG group, \*\**p*<0.01 vs HG group, \*\*\**p*<0.001 vs HG group, **** *p*<0.0001 vs HG group; ^#^*p*<0.05 vs HG+LG group, ^##^*p*<0.01 vs HG+LG group, ^###^*p*<0.001 vs HG+LG group, ^####^*p*<0.0001 vs HG+LG group. AGEs: advanced glycation end products; RAGE: receptor for advanced glycation end products.

## DISCUSSION

Our study reveals the role of ZBP1-mediated PANoptosis in cognitive impairment induced by recurrent episodes of hypoglycemia in mice with type 2 diabetes. In the Graphical Abstract (Figure 6), we describe the mechanism of brain damage in mice with type 2 diabetes during hypoglycemic episodes. A sensor of the PANoptosome, ZBP1, was activated by hypoglycemia in the brain endothelial cell line bEnd3, thereby activating pyroptosis, apoptosis and necroptosis. PANoptosis of endothelial cells *in vitro* and hippocampal inflammation *in vivo* led to cognitive impairment in mice.

**Figure 6.**
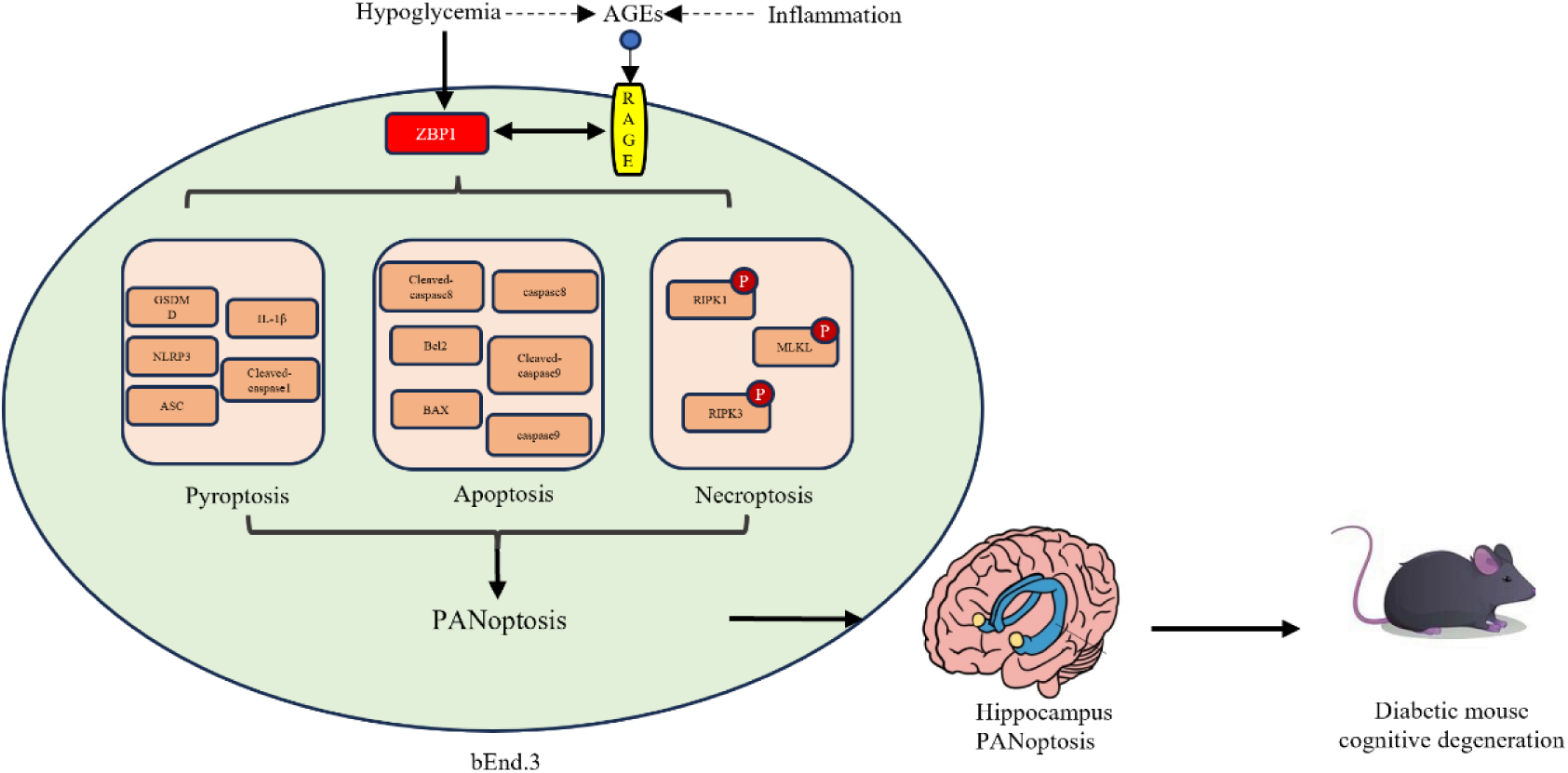
The graphical abstract of ZBP1-mediated PANoptosis activated by hypoglycemia contributes to cognitive degeneration in type 2 diabetes. In type 2 diabetic mouse brain, when hypoglycemia happens, the sensor of PANoptosome, ZBP1, was initially activated in the endothelial cells. After that, molecules involved in pyroptosis, apoptosis and necroptosis were overtly raised in bEnd.3 cell line. Moreover, the AGEs-RAGE axis was activated by hypoglycemia and ZBP1, which aggregated the ZBP1-mediated PANoptosis in bEnd.3. Based on PANoptosis was activated, endothelial cells injured and hippocampus was in an inflammatory state thus led to spatial learning and cognitive degeneration.

Previous studies have reported that hypoglycemia leads to cognitive impairment in patients with diabetes [31, 32]. However, another prospective study reported that moderate hypoglycemia did not increase the risk of cognitive impairment in middle-aged patients with abnormal blood glucose levels [33]. The brain is very sensitive to blood glucose fluctuations, and severe hypoglycemia causes irreversible damage to the mouse brain, resulting in slow responses, reduced learning and memory ability and chronic cognitive impairment. Therefore, it is important to study the compensatory mechanism of the brain after hypoglycemia and to find ways to block it. A previous study revealed that the activation of mitochondrial autophagy [34] alleviated cognitive degeneration caused by recurrent moderate hypoglycemia in diabetic mice. A recent study reported that lactic acid supplementation after hypoglycemia attenuated the cognitive dysfunction caused by recurrent moderate hypoglycemia in diabetic mice [35]. There have also been reports of the use of medications to prevent cognitive impairment caused by hypoglycemia [36]. Our study supports a strong association between hypoglycemia and cognitive impairment in patients with diabetes (Tables 1–2). Notably, this study focused on mice with type 2 diabetes, where episodes of hypoglycemia are more subtle than those in mice with type 1 diabetes [37]. Nevertheless, cognitive impairment is induced by hypoglycemia, so it is necessary and novel to study models of type 2 diabetes as well.

As “first-line barriers” that regulate blood flow and vasodilation function, endothelial cells are closely related to cognitive impairment [38]. Yaping Lu et al. reported that endothelial TFEB signaling-mediated autophagy disorders lead to microglial activation and cognitive dysfunction [39]. Moreover, markers of vascular cognitive impairment and dementia are summarized in a review [40]. Acute inflammation caused by infection and other factors can damage the endothelium and the blood‒brain barrier [41]; noninfectious factors such as hypertension [42], diabetes [43] and other chronic injuries can cause endothelial inflammation and cognitive degradation as well. However, the specific mechanisms by which these factors lead to endothelial inflammation differ. This study further revealed the role of inflammatory PCD-PANoptosis in the inflammatory injury of endothelial cells induced by hypoglycemia, providing new insights for future research and targeted therapy.

To date, widespread apoptosis related to diabetes is involved mainly in complications such as diabetic liver injury [44] and diabetic kidney injury [19]. Moreover, the liver can be targeted to improve systemic glucose homeostasis in T2DM rats [45]. *Song* and colleagues reported that Roux-en-Y gastric bypass surgery regulates the expression of TFF3 in the liver of ZDF rats, thereby activating the PI3K/Akt pathway and improving T2DM. Moreover, as the main organ for sugar and lipid metabolism, the liver plays a crucial role in regulating metabolism during the progression of diabetes. However, little is known about the mechanisms of PANoptosis related to the cognitive impairment caused by hypoglycemia, a complication of diabetes treatment. In this study, we further revealed that the activation of PANoptosomes in brain endothelial cells is mediated by the ZBP1 sensor, suggesting that ZBP1 or blocking intracerebral inflammation may be potential targets for treatment. In contrast, a previous study revealed that following sciatic nerve injury in a streptozotocin rodent model of type I diabetes, the mobilization of RNAs into injured axons was attenuated and correlated with decreased axonal regeneration. This failure of axonal RNA localization resulted from decreased levels of the RNA binding protein ZBP1 [46]. The different actions of ZBP1 may be related to the use of different animal models, the different target organs and the different degrees of inflammatory damage. However, the role of ZBP1 in hypoglycemia-related diseases has not been reported, and the specific mechanism remains unknown.

AGEs-RAGE have long been known to be involved in the pathology of pulmonary diseases [47]. Recent studies have shown the pivotal role of the AGE-RAGE axis in brain aging [48]. The authors reported that upon binding to the signal transduction receptor RAGE, AGEs can initiate proinflammatory pathways and exacerbate oxidative stress and neuroinflammation, thus impairing neuronal function and cognition. AGE-RAGE signaling induces programmed cell death, disrupts the blood–brain barrier and promotes protein aggregation, further compromising brain health. This finding is in line with the findings of our study, which revealed a proinflammatory effect of ZBP1-mediated PANoptosis induced by hypoglycemia in the hippocampus of mice with type 2 diabetes, which initiated AGE-RAGE signaling in brain endothelial cells. A reduction in AGE deposition in brain tissue through novel pharmacological therapeutics, such as ZBP1 knockdown, shows great promise for mitigating the cognitive decline associated with brain aging. Recently, a study revealed an association between AGE-RAGE signaling and apoptosis in cerebral ischemia‒reperfusion [49], which suggested that inhibiting apoptosis by regulating the AGE-RAGE axis was an effective way to attenuate cerebral injury. Moreover, studies have suggested the role of RAGE inhibitors in neurodegenerative diseases[50]. Our study provides a novel direction for exploring neural degenerative diseases.

A limitation of this study is that the mechanisms underlying ZBP1 activation by hypoglycemia and ZBP1 induction of PANoptosis in endothelial cells has not been revealed. The innovation of this study is that ZBP1-mediated PANoptosis has been reported for the first time in hypoglycemia-induced cognitive impairment in mice with type 2 diabetes. Differences between animal models of hypoglycemia-related diabetes and human patients should be noted. Hypoglycemia occurs randomly and inconsistently in diabetic patients[51]. However, animal models of hypoglycemic diabetic mice differ across studies [24, 52]. Our study was based on a previous study[24] and demonstrated a degree of cognitive impairment in type 2 diabetic mice. In the future, we will continue to study the potential mechanism involved in the interactions between AGER and ZBP1-mediated PANoptosomes to provide a new direction and solid evidence for translational research on clinical prevention and targets for treatment of cognitive impairment caused by hypoglycemia in diabetic patients.

## Authors Contribution

Conceptualization and study design: LN, LWP; animal procedures: LN; histology, membrane fractionation, immunoblotting, and data analysis: LWP, HA,CJX; manuscript draft: LN; funding acquisition and project administration: LSX. All authors: manuscript editing, review, and final approval.

## Availability of data and materials

The datasets used or analysed during the current study are available from the corresponding author on reasonable request. The associated accession number of RNA sequencing database is PRJCA046743 in BIG Submission (https://ngdc.cncb.ac.cn/gsub/submit/gsa/subCRA049776/finishedOverview; China National Center for Bioinformation).

## Ethics approval and consent to participate

The Laboratory Animal Management Committee of Chongqing Medical University approved this study. The animal treatment method fully complied with the relevant guidelines of the Animal Protection Law of the People’s Republic of China and obtained approval from the Institutional Ethics Committee of Chongqing Medical University of the State Science and Technology Commission of the People’s Republic of China (Approval number: IACUC-CQMU-2023-0065).

## Patient consent for publication

None.

## Acknowledgements

Not applicable.

---

**Supplementary Figure 1.**
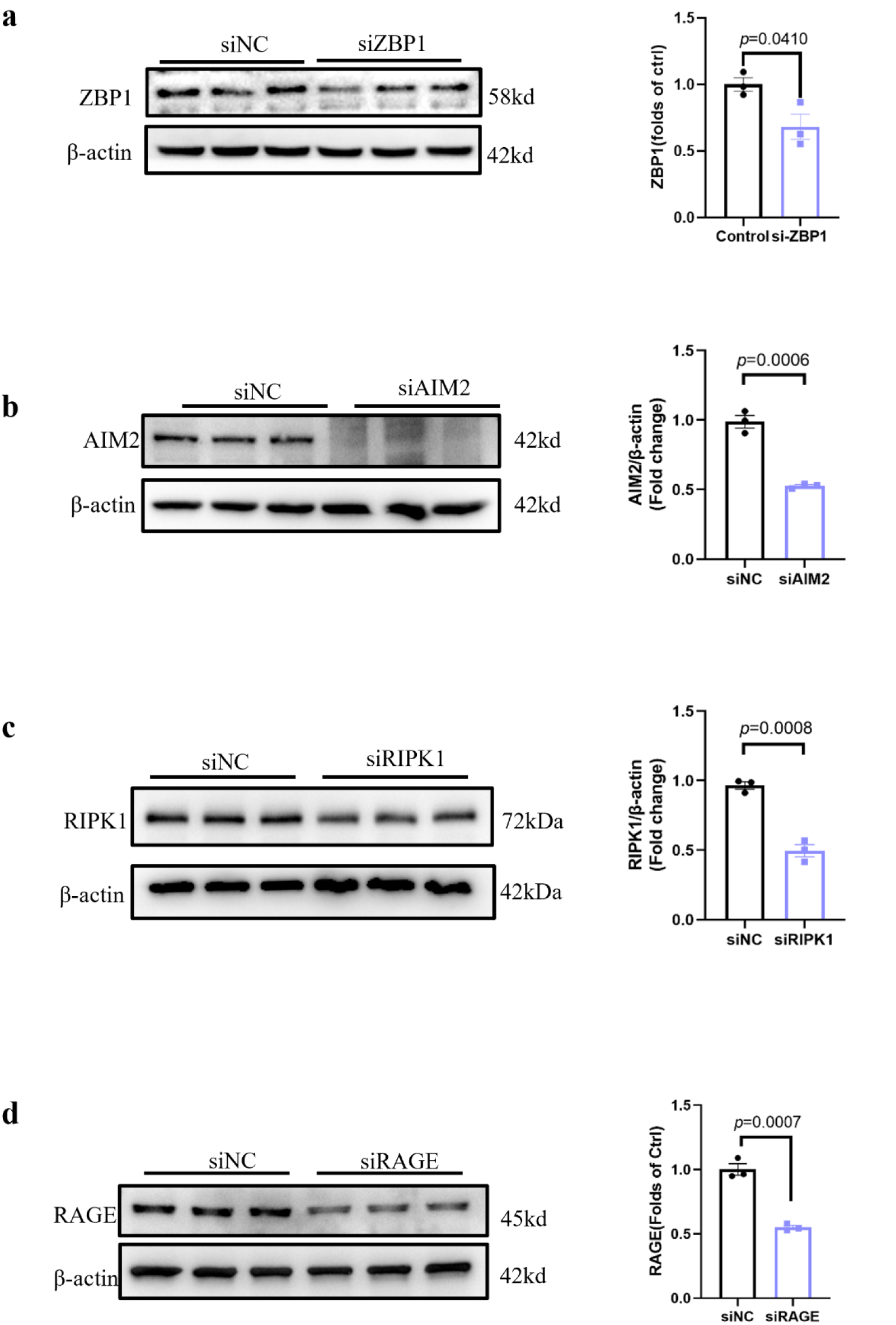
Validation of the successful knockdown of the protein expression of ZBP1(a), AIM2(b), RIPK1(c) and RAGE(d) in bEnd.3.

## References

1. Kaur, J., E. R. Seaquist 2023 Hypoglycaemia in type 1 diabetes mellitus: risks and practical prevention strategies. Nat Rev Endocrinol 19(3): p. 177–186. 10.1038/s41574-022-00762-8

2. International Hypoglycaemia Study, Group 2019 Hypoglycaemia, cardiovascular disease, and mortality in diabetes: epidemiology, pathogenesis, and management. Lancet Diabetes Endocrinol 7(5): p. 385–396. 10.1016/S2213-8587(18)30315-2

3. Verhulst, C. E. M., T. W. Fabricius, G. Nefs, R. P. C. Kessels, F. Pouwer, et al. 2022 Consistent Effects of Hypoglycemia on Cognitive Function in People With or Without Diabetes. Diabetes Care 45(9): p. 2103–2110. 10.2337/dc21-2502

4. Galvez, T., M. Lotierzo, G. Conquet, Q. Verani, C. Aguilhon, et al. 2023 Factitious hypoglycemia in insulin-treated diabetic patients. Ann Endocrinol (Paris) 84(3): p. 364–366. 10.1016/j.ando.2023.03.015

5. Sandoval, D. A., M. E. Patti 2023 Glucose metabolism after bariatric surgery: implications for T2DM remission and hypoglycaemia. Nat Rev Endocrinol 19(3): p. 164–176. 10.1038/s41574-022-00757-5

6. Zhou, Y., L. Huang, W. Zheng, J. An, Z. Zhan, et al. 2018 Recurrent nonsevere hypoglycemia exacerbates imbalance of mitochondrial homeostasis leading to synapse injury and cognitive deficit in diabetes. Am J Physiol Endocrinol Metab 315(5): p. E973–E986. 10.1152/ajpendo.00133.2018

7. McNay, E. C., V. E. Cotero 2010 Mini-review: impact of recurrent hypoglycemia on cognitive and brain function. Physiol Behav 100(3): p. 234–8. 10.1016/j.physbeh.2010.01.004

8. Gao, R., L. Ren, Y. Zhou, L. Wang, Y. Xie, et al. 2021 Recurrent non-severe hypoglycemia aggravates cognitive decline in diabetes and induces mitochondrial dysfunction in cultured astrocytes. Mol Cell Endocrinol 526(111192. 10.1016/j.mce.2021.111192

9. Zhang, X. G., Y. Q. Zhang, D. K. Zhao, J. X. Wu, J. Zhao, et al. 2014 Relationship between blood glucose fluctuation and macrovascular endothelial dysfunction in type 2 diabetic patients with coronary heart disease. Eur Rev Med Pharmacol Sci 18(23): p. 3593–600.

10. Chen, J., L. Lippo, R. Labella, S. L. Tan, B. D. Marsden, et al. 2021 Decreased blood vessel density and endothelial cell subset dynamics during ageing of the endocrine system. EMBO J 40(1): p. e105242. 10.15252/embj.2020105242

11. Ge, D., T. Luo, Y. Sun, M. Liu, Y. Lyu, et al. 2024 Natural diterpenoid EKO activates deubiqutinase ATXN3 to preserve vascular endothelial integrity and alleviate diabetic retinopathy through c-fos/focal adhesion axis. Int J Biol Macromol 260(Pt 2): p. 129341. 10.1016/j.ijbiomac.2024.129341

12. Zhang, J., D. Zhang, J. S. McQuade, M. Behbehani, J. Z. Tsien, et al. 2002 c-fos regulates neuronal excitability and survival. Nat Genet 30(4): p. 416–20. 10.1038/ng859

13. El Mahmoudi, N., C. Laurent, D. Pericat, I. Watabe, A. Lapotre, et al. 2023 Long-lasting spatial memory deficits and impaired hippocampal plasticity following unilateral vestibular loss. Prog Neurobiol 223(102403. 10.1016/j.pneurobio.2023.102403

14. Won, S. J., B. H. Yoo, T. M. Kauppinen, B. Y. Choi, J. H. Kim, et al. 2012 Recurrent/moderate hypoglycemia induces hippocampal dendritic injury, microglial activation, and cognitive impairment in diabetic rats. J Neuroinflammation 9(182. 10.1186/1742-2094-9-182

15. Kealy, J., C. Murray, E. W. Griffin, A. B. Lopez-Rodriguez, D. Healy, et al. 2020 Acute Inflammation Alters Brain Energy Metabolism in Mice and Humans: Role in Suppressed Spontaneous Activity, Impaired Cognition, and Delirium. J Neurosci 40(29): p. 5681–5696. 10.1523/JNEUROSCI.2876-19.2020

16. Sun, X., Y. Yang, X. Meng, J. Li, X. Liu, et al. 2024 PANoptosis: Mechanisms, biology, and role in disease. Immunol Rev 321(1): p. 246–262. 10.1111/imr.13279

17. Zheng, M., T. D. Kanneganti 2020 The regulation of the ZBP1-NLRP3 inflammasome and its implications in pyroptosis, apoptosis, and necroptosis (PANoptosis). Immunol Rev 297(1): p. 26–38. 10.1111/imr.12909

18. Lee, S., R. Karki, Y. Wang, L. N. Nguyen, R. C. Kalathur, et al. 2021 AIM2 forms a complex with pyrin and ZBP1 to drive PANoptosis and host defence. Nature 597(7876): p. 415–419. 10.1038/s41586-021-03875-8

19. Han, G., K. Hu, T. Luo, W. Wang, D. Zhang, et al. 2025 Research progress of non-coding RNA regulating the role of PANoptosis in diabetes mellitus and its complications. Apoptosis 30(3-4): p. 516–536. 10.1007/s10495-024-02066-w

20. Chang, X., B. Wang, Y. Zhao, B. Deng, P. Liu, et al. 2024 The role of IFI16 in regulating PANoptosis and implication in heart diseases. Cell Death Discov 10(1): p. 204. 10.1038/s41420-024-01978-5

21. Lv, Z., J. Hu, H. Su, Q. Yu, Y. Lang, et al. 2025 TRAIL induces podocyte PANoptosis via death receptor 5 in diabetic kidney disease. Kidney Int 107(2): p. 317–331. 10.1016/j.kint.2024.10.026

22. Luo, M., Y. Hu, D. Lv, L. Xie, S. Yang, et al. 2022 Recurrent Hypoglycemia Impaired Vascular Function in Advanced T2DM Rats by Inducing Pyroptosis. Oxid Med Cell Longev 2022(7812407. 10.1155/2022/7812407

23. Yu, Q., D. Yang, B. Ran, J. Pan, Y. Song, et al. 2025 SIRT4-Mediated Deacetylation of PRDX3 Attenuates Liver Ischemia Reperfusion Injury by Suppressing Ferroptosis. Int J Biol Sci 21(10): p. 4663–4682. 10.7150/ijbs.114510

24. Zuo, D., Y. Peng, G. Zhao, Z. Cheng, M. Luo, et al. 2025 Hypoglycemia Induces Diabetic Macrovascular Endothelial Dysfunction via Endothelial Cell PANoptosis, Macrophage Polarization, and VSMC Fibrosis. Adv Sci (Weinh) e14530. 10.1002/advs.202414530

25. Tataei, A., B. Rahimi, H. L. Afshar, V. Alinejad, H. Jafarizadeh, et al. 2023 The effects of electronic nursing handover on patient safety in the general (non-COVID-19) and COVID-19 intensive care units: a quasi-experimental study. BMC Health Serv Res 23(1): p. 527. 10.1186/s12913-023-09502-8

26. Lissner, L. J., K. M. Wartchow, A. P. Toniazzo, C. A. Goncalves, L. Rodrigues 2021 Object recognition and Morris water maze to detect cognitive impairment from mild hippocampal damage in rats: A reflection based on the literature and experience. Pharmacol Biochem Behav 210(173273. 10.1016/j.pbb.2021.173273

27. McLennan, S. N., A. K. Lam, J. L. Mathias, S. A. Koblar, M. A. Hamilton-Bruce, et al. 2011 Role of vasodilation in cognitive impairment. Int J Stroke 6(3): p. 280. 10.1111/j.1747-4949.2011.00601.x

28. Lisman, J., G. Buzsaki, H. Eichenbaum, L. Nadel, C. Ranganath, et al. 2017 Viewpoints: how the hippocampus contributes to memory, navigation and cognition. Nat Neurosci 20(11): p. 1434–1447. 10.1038/nn.4661

29. McNeilly, A. D., J. R. Gallagher, A. T. Dinkova-Kostova, J. D. Hayes, J. Sharkey, et al. 2016 Nrf2-Mediated Neuroprotection Against Recurrent Hypoglycemia Is Insufficient to Prevent Cognitive Impairment in a Rodent Model of Type 1 Diabetes. Diabetes 65(10): p. 3151–60. 10.2337/db15-1653

30. Clyne, A. M. 2021 Endothelial response to glucose: dysfunction, metabolism, and transport. Biochem Soc Trans 49(1): p. 313–325. 10.1042/BST20200611

31. Lin, L., Y. Wu, Z. Chen, L. Huang, L. Wang, et al. 2021 Severe Hypoglycemia Contributing to Cognitive Dysfunction in Diabetic Mice Is Associated With Pericyte and Blood-Brain Barrier Dysfunction. Front Aging Neurosci 13(775244. 10.3389/fnagi.2021.775244

32. Sheen, Y. J., W. H. Sheu 2016 Association between hypoglycemia and dementia in patients with type 2 diabetes. Diabetes Res Clin Pract 116(279–87. 10.1016/j.diabres.2016.04.004

33. Cukierman-Yaffe, T., J. Bosch, H. Jung, Z. Punthakee, H. C. Gerstein 2019 Hypoglycemia and Incident Cognitive Dysfunction: A Post Hoc Analysis From the ORIGIN Trial. Diabetes Care 42(1): p. 142–147. 10.2337/dc18-0690

34. Wu, K., C. Huang, W. Zheng, Y. Wu, Q. Huang, et al. 2024 Activation of mitophagy improves cognitive dysfunction in diabetic mice with recurrent non-severe hypoglycemia. Mol Cell Endocrinol 580(112109. 10.1016/j.mce.2023.112109

35. Wu, Y., R. Gao, Q. Huang, C. Huang, L. Wang, et al. 2025 Lactate supplementation after hypoglycemia alleviates cognitive dysfunction induced by recurrent non-severe hypoglycemia in diabetic mice. Exp Neurol 383(115037. 10.1016/j.expneurol.2024.115037

36. Jackson, D. A., T. Michael, A. Vieira de Abreu, R. Agrawal, M. Bortolato, et al. 2018 Prevention of Severe Hypoglycemia-Induced Brain Damage and Cognitive Impairment With Verapamil. Diabetes 67(10): p. 2107–2112. 10.2337/db18-0008

37. van Mark, G., S. Lanzinger, S. Sziegoleit, F. J. Putz, M. Durmaz, et al. 2019 Characteristics of Patients with Type-1 or Type-2 Diabetes Receiving Insulin Glargine U300: An Analysis of 7268 Patients Based on the DPV and DIVE Registries. Adv Ther 36(7): p. 1628–1641. 10.1007/s12325-019-00983-w

38. Fang, Y. C., Y. C. Hsieh, C. J. Hu, Y. K. Tu 2023 Endothelial Dysfunction in Neurodegenerative Diseases. Int J Mol Sci 24(3): p. 10.3390/ijms24032909

39. Lu, Y., X. Chen, X. Liu, Y. Shi, Z. Wei, et al. 2023 Endothelial TFEB signaling-mediated autophagic disturbance initiates microglial activation and cognitive dysfunction. Autophagy 19(6): p. 1803–1820. 10.1080/15548627.2022.2162244

40. Hosoki, S., G. K. Hansra, T. Jayasena, A. Poljak, K. A. Mather, et al. 2023 Molecular biomarkers for vascular cognitive impairment and dementia. Nat Rev Neurol 19(12): p. 737–753. 10.1038/s41582-023-00884-1

41. Greene, C., R. Connolly, D. Brennan, A. Laffan, E. O’Keeffe, et al. 2024 Blood-brain barrier disruption and sustained systemic inflammation in individuals with long COVID-associated cognitive impairment. Nat Neurosci 27(3): p. 421–432. 10.1038/s41593-024-01576-9

42. Uiterwijk, R., M. Huijts, J. Staals, R. P. Rouhl, P. W. De Leeuw, et al. 2016 Endothelial Activation Is Associated With Cognitive Performance in Patients With Hypertension. Am J Hypertens 29(4): p. 464–9. 10.1093/ajh/hpv122

43. Phoenix, A., R. Chandran, A. Ergul 2022 Cerebral Microvascular Senescence and Inflammation in Diabetes. Front Physiol 13(864758. 10.3389/fphys.2022.864758

44. Neira, G., S. Becerril, V. Valenti, R. Moncada, V. Catalan, et al. 2024 FNDC4 reduces hepatocyte inflammatory cell death via AMPKalpha in metabolic dysfunction-associated steatotic liver disease. Clin Nutr 43(9): p. 2221–2233. 10.1016/j.clnu.2024.08.007

45. Song, K., X. Kong, Y. Xian, Z. Yu, M. He, et al. 2025 Roux-en-Y gastric bypass improves liver and glucose homeostasis in Zucker diabetic fatty rats by upregulating hepatic trefoil factor family 3 and activating the phosphatidylinositol 3-kinase/protein kinase B pathway. Surg Obes Relat Dis 21(7): p. 792–805. 10.1016/j.soard.2024.12.024

46. Jones, J. I., C. J. Costa, C. Cooney, D. C. Goldberg, M. Ponticiello, et al. 2021 Failure to Upregulate the RNA Binding Protein ZBP After Injury Leads to Impaired Regeneration in a Rodent Model of Diabetic Peripheral Neuropathy. Front Mol Neurosci 14(728163. 10.3389/fnmol.2021.728163

47. Sharma, A., S. Kaur, M. Sarkar, B. C. Sarin, H. Changotra 2021 The AGE-RAGE Axis and RAGE Genetics in Chronic Obstructive Pulmonary Disease. Clin Rev Allergy Immunol 60(2): p. 244–258. 10.1007/s12016-020-08815-4

48. Vitorakis, N., C. Piperi 2024 Pivotal role of AGE-RAGE axis in brain aging with current interventions. Ageing Res Rev 100(102429. 10.1016/j.arr.2024.102429

49. Wang, W., W. Zhao, X. Song, H. Wang, L. Gu 2025 Zhongfeng decoction attenuates cerebral ischemia-reperfusion injury by inhibiting autophagy via regulating the AGE-RAGE signaling pathway. J Ethnopharmacol 336(118718. 10.1016/j.jep.2024.118718

50. Reddy, V. P., P. Aryal, P. Soni 2023 RAGE Inhibitors in Neurodegenerative Diseases. Biomedicines 11(4): p. 10.3390/biomedicines11041131

51. Umpierrez, G., M. Korytkowski 2016 Diabetic emergencies - ketoacidosis, hyperglycaemic hyperosmolar state and hypoglycaemia. Nat Rev Endocrinol 12(4): p. 222–32. 10.1038/nrendo.2016.15

52. Guo, C., Y. Niu, X. Pan, D. Sharma, E. Lau, et al. 2025 Hypoglycemia promotes inner blood-retinal barrier breakdown and retinal vascular leakage in diabetic mice. Sci Transl Med 17(796): p. eadq5355. 10.1126/scitranslmed.adq5355

